# Supplementary motor area disinhibition during motor sequence learning: A TMS-EEG study

**DOI:** 10.1101/2024.02.26.581077

**Authors:** Sophie Thong, Elizabeth Doery, Mana Biabani, Nigel C. Rogasch, Trevor T. -J. Chong, Joshua Hendrikse, James P. Coxon

**Author notes:** Correspondence to: James P. Coxon School of Psychological Sciences 18 Innovation Walk, Monash University Victoria 3800, Australia. **Competing interests** T.T.-J.C has received honoraria for lectures from Roche. The remaining authors declare no competing interests.

## Abstract

**Background:** In primary motor cortex, changes in excitatory and inhibitory neurotransmission (E:I balance) accompany motor sequence learning. In particular, there is an early reduction in inhibition (i.e., disinhibition). The supplementary motor area (SMA) is a key brain region involved in the learning of sequences, however the neurophysiological mechanisms within SMA which support motor sequence learning remain poorly understood. Disinhibition may also occur in SMA, but this possibility remains unexamined.

**Objective:** We investigated disinhibition within SMA during motor sequence learning using combined transcranial magnetic stimulation (TMS) and electroencephalography (EEG).

**Methods:** Twenty-nine healthy adults practiced a sequential motor task. TMS-evoked potentials (TEPs) resulting from SMA stimulation were measured with EEG before, during, and after practice. The N45 TEP peak was our primary measure of disinhibition. Furthermore, the slope of aperiodic EEG activity was included as an additional E:I balance measure.

**Results:** Significant improvements in task performance (i.e., learning) occurred with practice. We observed smaller N45 amplitudes during early learning relative to baseline (both *p* < .01), indicative of disinhibition. Intriguingly, aperiodic exponents increased as learning progressed and were associated with greater sequence learning (*p* < .05).

**Conclusion:** Our results show disinhibition within SMA during the planning phase of motor sequence learning and thus provide novel understanding on the neurophysiological mechanisms within higher-order motor cortex that accompany new sequence learning.

## Introduction

Our capacity to learn and execute sequential actions with increasing speed and accuracy is fundamental to the development of motor skills [1]. This process, referred to as motor sequence learning, is characterised by rapid improvements in execution during early stages of learning [1], and is facilitated by the ‘pre-planning’ of relevant actions prior to movement [2]. Motor planning and learning have been associated with changes in the balance between excitatory and inhibitory neurotransmission (E:I balance) within primary motor cortex (M1) [3-5]. The Supplementary Motor Area (SMA) is known to play a central role in sequencing [6, 7], however the neurophysiological mechanisms in SMA that support motor sequence learning remain poorly understood.

Disinhibition, i.e., reduced gamma-aminobutyric acid (GABA) transmission, is a prerequisite for the neuroplasticity processes supporting learning [8, 9], and has been demonstrated to occur within M1 during motor sequence learning [3, 4, 10]. Furthermore, the magnitude of M1 disinhibition has been linked to motor sequence learning [11], although this is not consistently observed [4]. Interestingly, shifting the E:I balance towards excitation via electrical stimulation of SMA during movement preparation was found to improve action execution [12]. However, the occurrence and significance of disinhibition within SMA during motor sequence learning remains undetermined.

Local and distributed changes in E:I balance can be measured via combined use of transcranial magnetic stimulation (TMS) and electroencephalography (EEG), whereby TMS is administered to a cortical site and simultaneous EEG recordings register TMS-evoked potentials (TEPs) [13, 14]. A TEP peak occurring approximately 45ms after M1 stimulation (i.e., the N45) has been linked to GABA_a_ [15] and N-methyl-D-aspartate (NMDA) receptor signalling [16]. Based on these findings, smaller N45 amplitudes reflect an E:I balance shift towards disinhibition [15, 16]. In addition to TEPs, the slope of aperiodic EEG activity within the frequency domain is sensitive to changes in E:I balance [17], with flatter (smaller) slopes suggesting reduced inhibition [18, 19]. Together, the N45 amplitude and aperiodic exponent may offer unique insights into the E:I dynamics supporting motor sequence learning.

We examined disinhibition in SMA and associated cortical regions during motor sequence learning using TMS-EEG. We hypothesised there would be an early reduction in N45 amplitude associated with learning, and that this would be associated with learning outcomes. As an additional E:I balance marker, we examined changes in the aperiodic exponent across learning, and anticipated a reduction in the exponent during learning. We acknowledge that a negative TEP peak occurring approximately 100ms after M1 stimulation (i.e., the N100), has been linked to GABAergic signalling [15] and cognition [20]. However, this peak may predominantly reflect the sensory effects of stimulation [21, 22]. Therefore, for completeness, we also report the N100, noting that this should be interpreted with caution.

## Methods

### Participants

Thirty-five adults without history of neurological illnesses were recruited through convenience sampling. All participants were screened for their eligibility to undergo TMS. Five participants were excluded due to noisy EEG data, while one participant was excluded for failing to complete the task as instructed. The total sample thus comprised 29 adults (14 female, *M*_age_ = 24.79 years, *SD*_age_ = 7.12). The final sample comprised 28 right-handed individuals and one ambidextrous individual as determined by the Edinburgh Handedness Inventory [23]. Ethical approval was obtained from the Monash University Human Research Ethics Committee.

### Design

This study utilised a repeated-measures, within-subjects design. Participants completed a three-hour session in which TMS-EEG data was recorded, with measures obtained at rest before and after the task, and during sequential motor task practice.

### Procedure

#### Administration of TMS and EEG

Figure 1A outlines the experimental protocol. Each session began by fitting the participant with a 64-electrode actiCap with slim electrodes in the 10-20 montage (Brain Products, Germany). Resting-motor threshold (RMT) was determined using surface electromyography (Ag/Ag-Cl electrodes) recorded from the right first dorsal interosseous (FDI) muscle. Single, monophasic TMS pulses were administered using a Magstim BiStim^2^ system with a figure-of-eight coil (70mm diameter) (The Magstim Company Ltd, UK). The motor ‘hotspot’ was determined as the cortical area that produced the largest and most consistent motor-evoked potential in the targeted FDI muscle. RMT was defined as the minimum stimulation intensity necessary to evoke a peak-to-peak amplitude of ≥ 50 μV in the resting FDI at least four out of eight consecutive trials.

The SMA stimulation site corresponded to the SMA-proper, 2.5cm anterior to *Cz*, and 0.5cm left of the midline [24]. The TMS coil handle was held pointing to the left, i.e., perpendicular to the midsagittal plane. EEG was recorded at a sampling rate of 25kHz using BrainVision software and an actiChamp amplifier (Brain Products, Germany), and referenced online to channel *Oz*. Impedances were kept below 5kΩ.

For TEP acquisition, single-pulse TMS was applied at 110% of the participant’s RMT. During resting measures, TMS pulses were administered at variable intervals (7.5±2s) while participants viewed a fixation cross on a computer screen. Seventy TMS pulses were delivered to SMA, followed by 70 TMS pulses to the acromion process of the scapula (i.e., the bony prominence of the shoulder). This control site was included to measure peripherally-evoked potentials without cortical stimulation [22]) (Figure 1A). During each task block, TMS was administered pseudo-randomly on 22 out of 32 trials. The TMS pulse occurred during the planning phase, 750ms following target stimulus onset and 1250ms prior to motor execution (see Figure 1). Throughout the protocol, participants were supported by a chin rest and listened to white noise through earphones to mask the sound of TMS [14] (Figure 1B).

**Figure 1.**
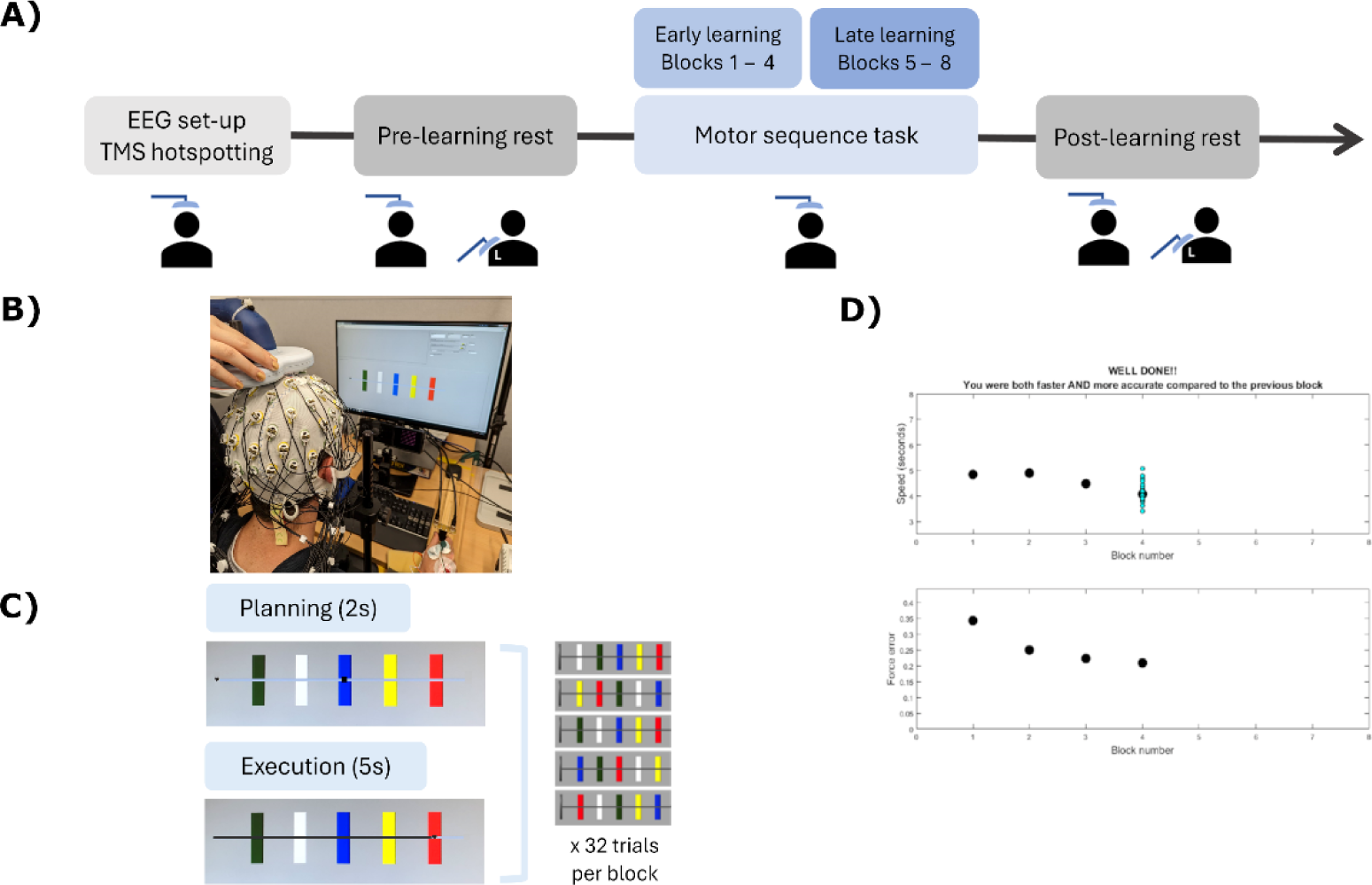
**A)** Timeline of the protocol. **B)** Experimental set-up. **C)** Schematic of all trials and the five motor execution sequences learnt throughout the task. **D)** An example of the feedback displayed following completion of a block. Uppermost panel: average trial duration (speed) per block. Second panel: force error relative to each target position.

### Sequential Visual Isometric Pinch Task (SVIPT)

Participants completed the SVIPT to assess motor sequence learning (Figure 1B) [25]. Participants controlled a cursor displayed on the screen by pinching a force transducer between their right index finger and thumb. On each trial, participants aimed to move the cursor to a series of coloured targets in a pre-specified order (red-blue-green-yellow-white) (Figure 1B). Participants were instructed to produce one force pulse (pinch) to reach each target, and then return to the ‘home’ position by relaxing their grip on the transducer. Cursor movement was proportional to the force registered at the transducer, whereby reaching the rightmost target required 45% of participants’ maximum voluntary contraction [3]. Participants completed three practice trials in a simple ascending/descending order to gain familiarity with the protocol, and then completed eight task blocks of 32 trials each.

Each trial involved a two second planning and five second execution phase. Coloured targets were displayed during the planning phase (Figure 1C). Participants were instructed to commence execution once a small black square aligned with the middle target disappeared. Across trials, target positions varied pseudo-randomly between five motor sequence execution orders (Figure 1C). Specifically, participants had to retrieve and execute a motor sequence on each trial, and learned all motor execution sequences concurrently throughout the session (i.e., interleaved motor practice). Participants were given feedback on their speed and accuracy after each block (Figure 1D) [26].

The primary outcome variable, skill, was calculated as a composite of speed and accuracy on each trial, and then averaged across trials for each condition (with TMS, without TMS). Skill was calculated for each participant using the following formula [26], where:

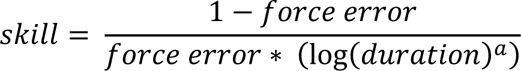

‘Force error’ refers to the summed distance between the cursor’s position at the peak of each force pulse and the target location (an inverse measure of accuracy). ‘Duration’ represents the time taken to complete a trial (s), i.e., execution speed. The parameter ‘a’ was set at 1.627 based on an independent dataset [3]. The skill measure used in analysis was the logarithm of this skill parameter, to ensure homogeneity of variance across participants [25]. Higher skill values reflect a shift in the speed-accuracy trade-off function towards faster and more accurate task performance, i.e., greater skill at performing the motor sequences.

### Data pre-processing

#### EEG data

Raw EEG data was processed in MATLAB (2017a) (The Mathworks Inc., US) using the TMS-EEG signal analyser (TESA) [27] and EEGLAB toolboxes [28].

#### TEP peak amplitudes

Rest and task data were cleaned following an established pipeline [27]. In brief, electrodes were first automatically rejected based on kurtosis. Rejected channels were interpolated in the final step of the pipeline. Data was epoched around each TMS pulse (i.e. from -1000ms to +1000ms), and baseline corrected (-500ms to -10ms). Data around the TMS pulse (-5ms to +10ms) was removed, followed by cubic data interpolation, and down-sampling to 1000Hz. Independent component analysis (ICA) was run to remove artifacts, involving automatic selection and visual inspection of components. Butterworth filters (1-100 Hz bandpass and 48-52 Hz bandstop) were then applied and data were re-referenced to the average of all channels.

Trials obtained during the SVIPT were averaged across Blocks 1-4 to encompass an ‘early learning’ phase, and across Blocks 5-8 to represent a ‘late learning’ phase. This resulted in 88 epochs for TEP averaging at each learning phase (less epochs rejected during pre-processing). The number of epochs for task TEPs was intentionally more than the rest timepoints (70 epochs), to ensure a similar number of epochs after data cleaning. On average, 85.13% of rest trials and 82.71% of task trials were retained.

#### Aperiodic exponents

Aperiodic exponent data were measured from the resting EEG between TMS pulses, epoched from 2500ms to 4500ms after each TMS pulse to avoid the evoked potential [14]) (70 trials). Aperiodic exponent data was also obtained from the two second planning phase of the SVIPT on trials without TMS administration (10 trials/block, 80 trials total across Blocks 1-8). For this analysis, data were downsampled to 1000 Hz, and high-pass filtered with 1 Hz cut-off. Electrodes were automatically rejected based on kurtosis (on average 4.2 electrodes removed across rest and task phases). Data were then interpolated, and re-referenced to the average of all channels. ICA was then run to remove artifacts.

Aperiodic exponents were extracted using Brainstorm (version 3.230621) [29]. The average power spectrum density was extracted for each participant at each timepoint (pre-learning rest, early learning, late learning, post-learning rest) using Welch’s method. The ‘Fitting-Oscillations-and-One-Over-F’ (FOOOF) spectral parametrisation model [17] was then applied to each participant’s average power spectrum density from 1-40 Hz, with peak widths ranging from 0.5 to 12 Hz, with no ‘knee’ component.

### Statistical analysis

Group-level statistical analyses were performed in JASP (version 0.17.2.1) [30] and MATLAB.

#### SVIPT motor learning data

A 2x8 repeated-measures ANOVA (rmANOVA) was conducted to assess skill learning with practice, and to examine whether the TMS pulse influenced learning. Factors were TMS (with-TMS, without-TMS) and Block (Blocks 1-8). Separate rmANOVAs were also conducted on the speed (Trial Duration) and accuracy (Force error components of the skill measure. Sphericity was assessed using Mauchly’s test, and Greenhouse-Geisser corrections were applied where ε < .75, while Huynh-Feldt corrections were used when ε > .75 [31].

For correlational analyses, early learning skill and late learning skill were quantified as skill averaged across Blocks 1-4 and Blocks 5-8 respectively, to align with TEP measures. Skill improvement was quantified as mean Block 8 skill - mean Block 1 skill (delta skill).

#### EEG data: TEP peak amplitudes

Cluster-based permutation t-tests implemented with the FieldTrip toolbox [32] were used to compare TEP peaks as a function of sequence learning, and stimulation condition (SMA vs. shoulder) (two-tailed α = .025; Monte Carlo, 10K permutations). For learning, data were compared across: 1) pre-learning rest vs. early learning, 2) early vs. late learning, 3) late learning vs. post-learning rest. Analyses included amplitudes from all electrodes averaged across 45-55ms (N45) and 85-140ms (N100) temporal windows [15]. For pre- and post-learning rest, visualisation of TEPs revealed that the N45 began around 35ms (Figure 3). Thus, the time window of N45 peaks recorded during rest was extended to 35-55ms.

For correlational analyses, we obtained mean TEP component amplitudes. Specifically, the amplitudes of electrodes in which significant differences were found between stimulation conditions and timepoints were first averaged across electrodes, and then time windows. We quantified changes in amplitude with learning as the difference in amplitude between time points (delta amplitude).

#### EEG data: Aperiodic exponents

Exponent values from electrodes (FOOOF model *R*^2^ > .8) where significant differences in the N45 were observed between pre-learning rest and early learning were included in group-level analyses. rmANOVA examined changes in exponents across the four timepoints. For correlational analyses, we calculated differences in mean exponents between timepoints (delta exponent). Positive delta values indicate I>E [17].

### Associations between neurophysiological data and learning metrics

To examine brain-behaviour relationships, we conducted correlations between learning measures (early skill, late skill, delta skill) and neurophysiological measures (baseline TEP component amplitude and exponent, and their delta change). We used Shapiro-Wilk tests to assess bivariate normality, and report Pearson’s or Spearman’s values where appropriate.

## Results

### Participants’ skill on the SVIPT task improved with practice

First, we confirmed that participants significantly improved in skill across SVIPT blocks (main effect of Block, *F* _27, 3.843_ = 62.199, *p* < .001, η^2^ = .617, Greenhouse-Geiser corrected) (Figure 2A). A main effect of TMS on participants’ skill was observed (*F* _27, 1_ = 4.246, *p* = .049, η^2^ = .003), although the TMS pulse during the planning phase only had a small effect on sequence execution (Figure 2B), and the TMS x Block interaction was not significant (*F* _27, 1_ = 1.781, *p* = .093, η^2^ = .006). Skill improvement was driven by improvements in speed and accuracy, with significant reductions in trial Force Error magnitude (*F* _27, 3.038_ = 27.166, *p* < .001, η^2^ = .502, Greenhouse-Geisser corrected), and Duration (s) (*F* _27, 3.230_ = 26.391, *p* < .001, η^2^ = .494, Greenhouse-Geisser corrected) across blocks (Figure 2C, D).

**Figure 2.**
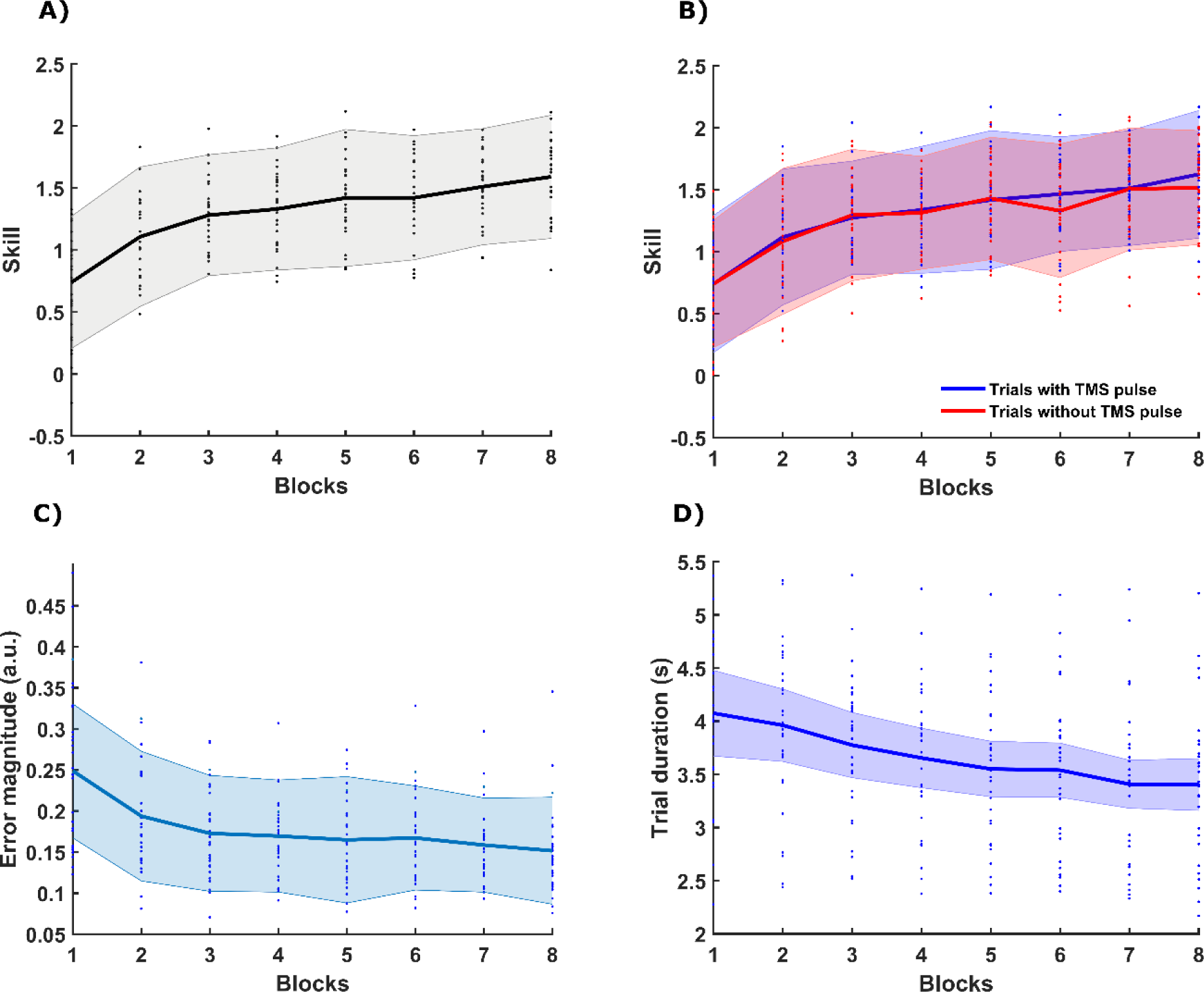
Participants improved significantly in skill (**A**), accuracy (**C**) and speed (**D**) (*ps* < .001) with practice. TMS pulses did not significantly alter participants’ performance (**B**). Shaded error bars represent ±SD.

### N45 amplitudes reflect effects of cortical stimulation

Next, we compared potentials evoked by SMA and shoulder stimulation. Relative to shoulder stimulation, SMA stimulation resulted in a prominent N45 (pre-learning rest, *p* < .001, post-learning rest, *p* = .002) and a smaller N100 (pre-learning, *p* = .019, post-learning, *p* <.001) (Figure 3B). Differences in the N45 were centred around the site of stimulation, while differences in the N100 were found at central-parietal electrodes (Figure 3A).

**Figure 3.**
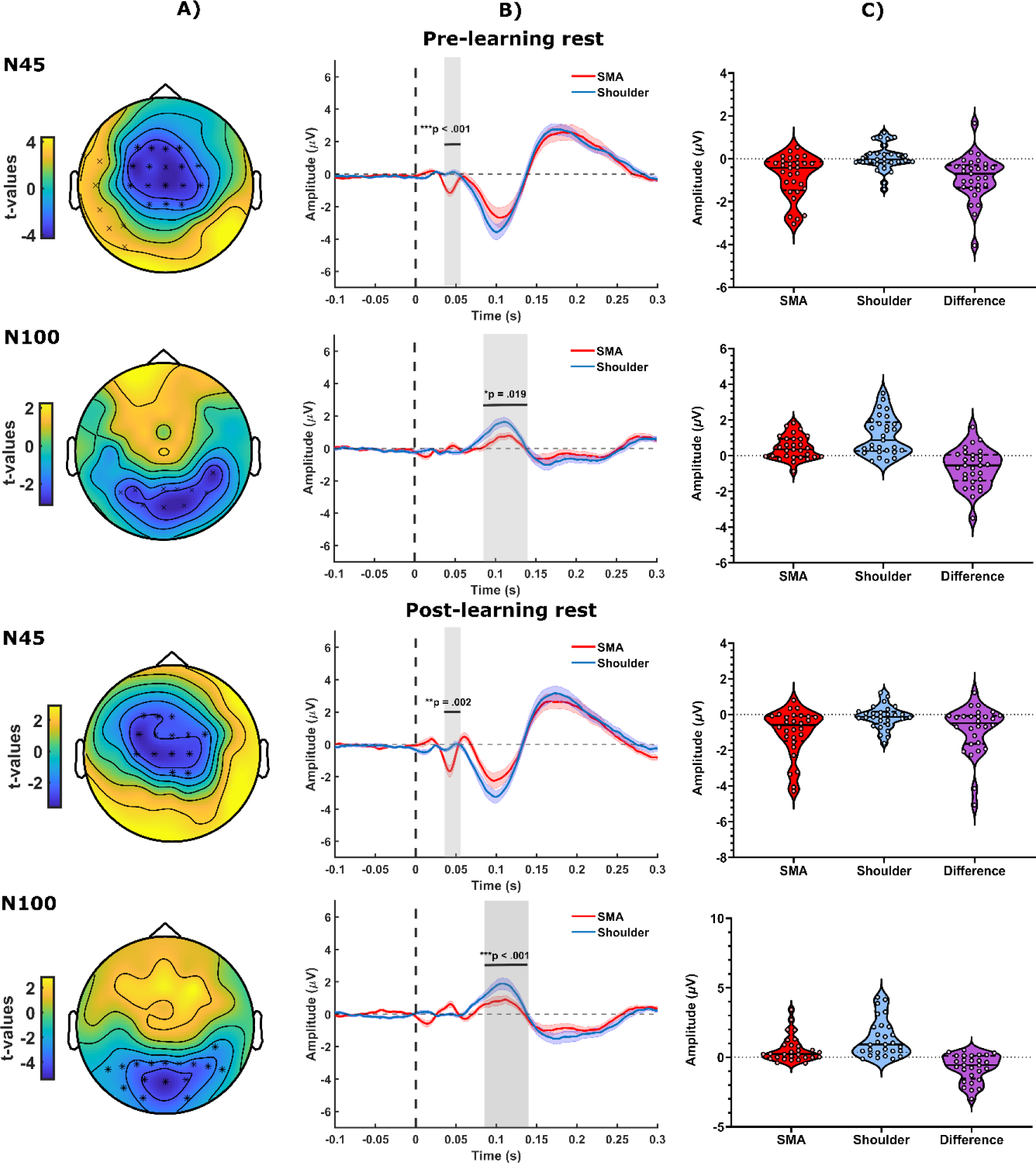
**A)** Topographical representation of TEP differences between SMA and shoulder stimulation. *Cluster *p* < .01, ^x^Cluster *p* < .025. Pre-learning negative cluster electrodes: *Fz, F1, F2, F4, FCz, FC1, FC2, FC4, C3, CPz, CP2, Cz, C2, C4* (N45); *Cz, C1, C2, C3, C5, CP1, CP3, CP6, Pz, P1, P2, P3* (N100). Post-learning cluster electrodes: *Fz, F1, F3, FCz*, *FC1, FC2, FC3, FC4, CPz, CP2, Cz, C1, C2, C3* (N45); *CP6, PO7, PO3, POz, PO4, Pz, P1, P2, P3, P4, P5, P6, P7, P8, O1, O2* (N100). **B)** Time course of TEPs averaged across cluster electrodes (dotted line at 0ms indicates TMS pulse). Shaded grey region and black horizontal lines depict averaged time windows where significant differences were observed. Coloured lines indicate group average for SMA stimulation (Blue) and Shoulder stimulation (Red), shaded error bars represent ±1SE. **C)** TEP amplitudes of each condition, and the difference between conditions (SMA amplitudes – Shoulder amplitudes). Negative difference values indicate a more negative N45 TEP amplitude following SMA stimulation, relative to shoulder stimulation.

We further probed the dissociation between potentials evoked by cortical vs. control stimulation by examining correlations between them. For the N45, there were no significant correlations between cortical and shoulder stimulation at either pre- or post-learning rest (*p*s > .08). For the N100, amplitudes correlated at post-learning rest (*p* = .013) (Supplementary Figure 1). Together, these results suggest the N45 reflects true effects of cortical stimulation, while the N100 evoked by SMA stimulation also reflects sensory effects of stimulation [22].

### N45 amplitude decreased from baseline to early learning

We examined changes in TEP peaks as a function of learning and observed a significant reduction in N45 amplitude at early learning relative to pre-learning rest across electrodes corresponding to the SMA (*p* = .0037) (Figure 4). Differences were also observed for the N100 across electrodes posterior to the stimulation site (*p* = .0072) (Figure 4). Our results provide evidence of disinhibition across the SMA and associated cortical regions during the planning phase of early motor sequence learning.

**Figure 4.**
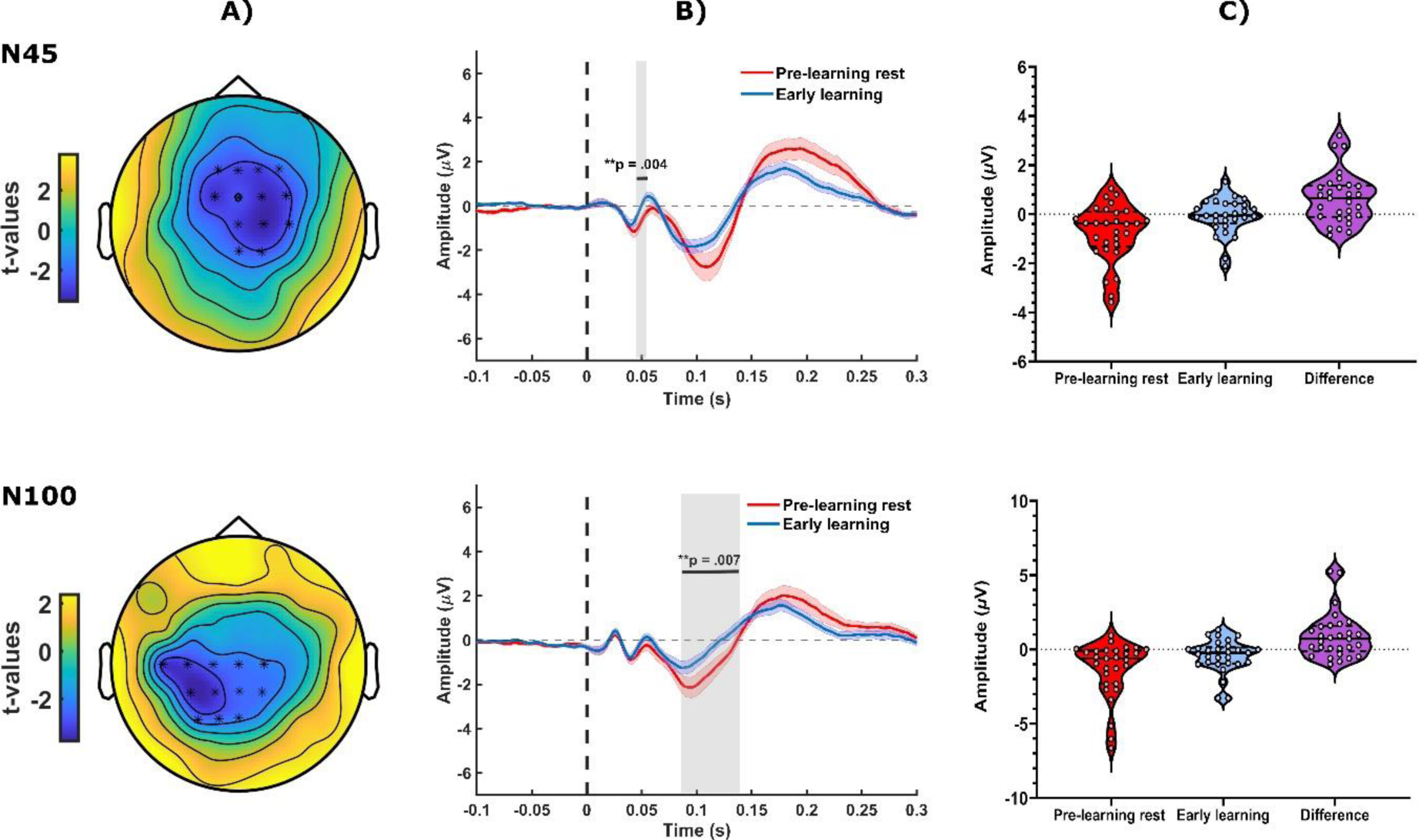
**A)** Topographical representation of TEP differences between pre-learning rest and early learning. *Cluster *p*-values < .01. Cluster electrodes: *Fz, F1, F2, F4, FCz, FC1, FC2, FC4, Cz, C2, C4, CPz, CP2* (N45); *Cz, C1, C2 C3, C5, CP1, CP2, CP3, Pz, P1, P3* (N100). **B)** Time course of TEPs averaged across cluster electrodes of both timepoints (dotted line at 0ms indicates TMS pulse). Shaded grey region and black horizontal lines depict averaged time windows where significant differences were observed. Coloured lines indicate group average for SMA stimulation during Early learning (Blue) and Pre-Learning Rest (Red), shaded error bars represent ±1SE. **C)** TEP amplitudes for each timepoint, and differences between them (early learning– pre-learning rest). Positive difference values indicate that early learning amplitudes are less negative (smaller) than pre-learning amplitudes.

### No change in TEP peak amplitudes across later stages of learning and rest

No significant differences in N45 amplitude were observed when comparing early learning to late learning, or late learning to post-learning rest (*p* > .025) (Figure 5).

**Figure 5.**
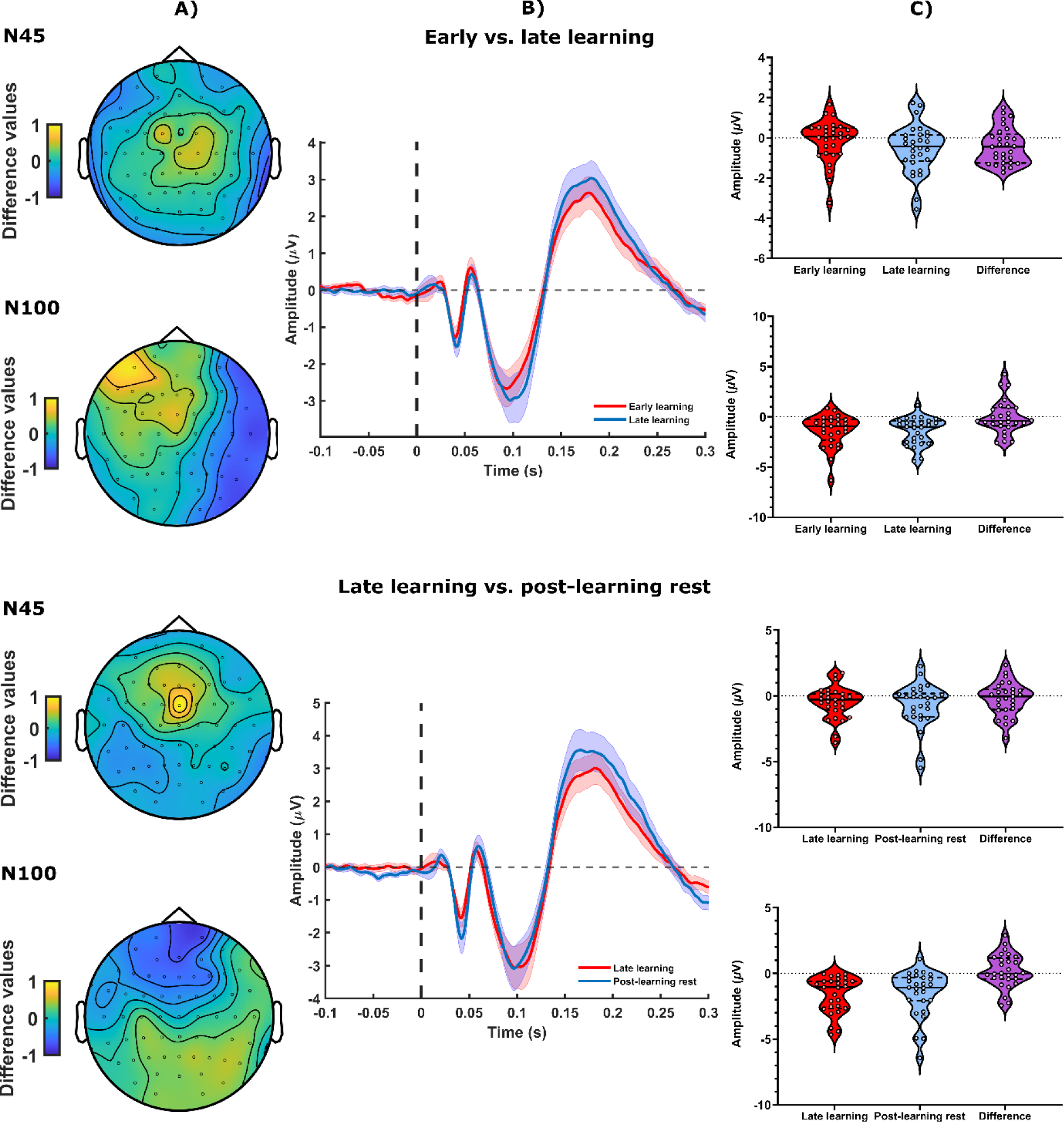
**A)** Topographical representation of differences in N45 and N100 amplitudes between conditions (no significant clusters). **B)** TEP amplitudes averaged across electrodes around the simulation site (*FC1, FC2, FCz, C1, C2* and *Cz*). Coloured lines indicate group average for SMA stimulation during Post-Learning Rest (Blue) and Late Learning (Red), shaded error bars represent ±1SE. **C)** TEP amplitudes of each timepoint, and differences between them. For visualisation purposes, outliers (Grubbs’ test) were removed for the N100 in the early vs. late condition (*N* = 2), and the N45 in the late vs. post-learning condition (*N* = 1).

### Relationships between TEP peak amplitudes and metrics of motor sequence learning

No significant associations were observed between pre-learning and early learning N45 amplitudes, and early, late or delta skill (all *p*s > .1). Similarly, delta N45 did not correlate with skill (all *p*s > .1). We observed significant correlations between N100 amplitudes and learning metrics (see Supplementary Data 2).

### Aperiodic exponents show a flatter slope after learning and correlates with skill improvements

Aperiodic exponents changed with time (*F*_27,2.774_ = 4.572, *p* = .007, η^2^ = .145, Huynh-Feldt corrected). Post-hoc analyses revealed a smaller exponent at post-learning rest relative to late learning (*t*_27_= 3.472, *p* = .005, Cohen’s *d* = .65, Bonferroni corrected) (Figure 6A). This change implies a post-learning shift in E:I balance, which differs from the early learning disinhibition reflected in TEPs. We thus explored correlations between delta amplitudes (early learning – pre-learning) and delta exponents (early learning – pre-learning; post-learning – late learning). No significant correlations were observed (all *p*s > .1), suggesting that the N45 and aperiodic exponent are independent markers of E:I balance.

**Figure 6.**
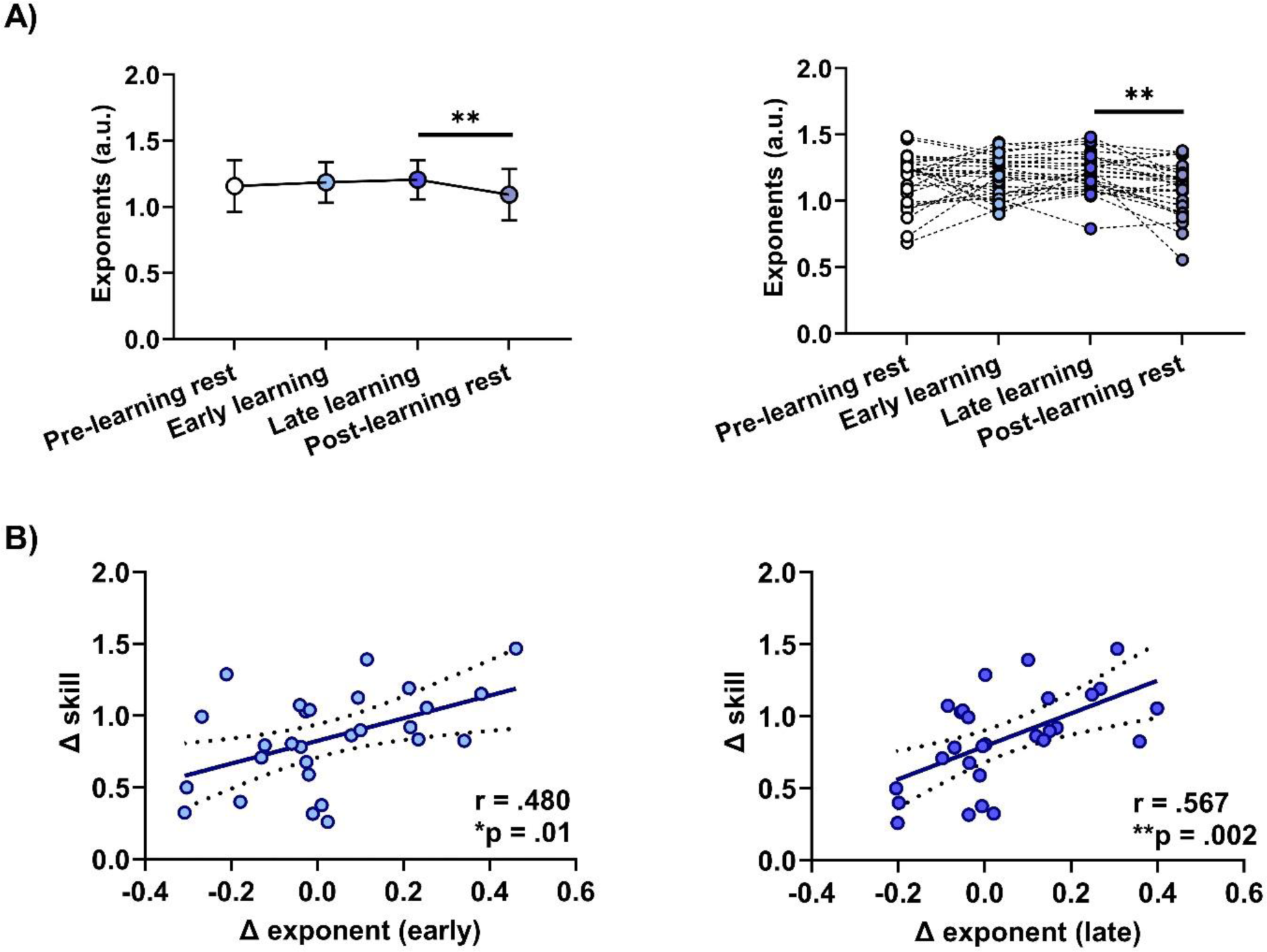
**A)** Aperiodic exponents over time. Left panel: mean exponents ± 1SD. Right panel: individual exponents. **B)** Correlations between skill improvements and changes in the aperiodic exponent from pre-learning rest to early learning (left panel) and late learning (right panel). The aperiodic exponent was averaged across the N45 cluster depicted in Figure 2. ** *p* = .005

We observed a marginally significant association between the pre-learning aperiodic exponent and early skill (ρ =.374, *p* = .047). We also found significant positive relationships between the delta exponent and delta skill, where an increase in the aperiodic exponent was associated with greater skill improvements (early delta: *r =* .480*, p =* .01, late delta: *r* = .567, *p* = .002) (Figure 6B).

## Discussion

We investigated disinhibition within SMA during motor sequence learning. Firstly, as hypothesised, we observed a reduction in the N45 during early learning, providing evidence of disinhibition across a cluster of electrodes centred over SMA. Secondly, we observed a smaller aperiodic exponent following learning, indicating a shift in E:I towards increased excitation and/or reduced inhibition. Interestingly, greater improvements in skill were associated with a task-related shift towards larger aperiodic exponents. Together, our findings implicate the modulation of inhibition in higher-order motor cortex as a mechanism supporting motor sequence learning.

### Changes in TEPs reveal disinhibition within SMA during early sequence learning

Functional magnetic resonance imaging studies in humans and single-cell recordings in primates demonstrate SMA activation as new action sequences are learned [6]. However, these techniques are limited by their low temporal resolution and spatial coverage, respectively. Here, we overcome these limitations via EEG recordings of TEPs following SMA stimulation. We demonstrate that N45 amplitude is reduced from baseline to early learning. TMS-EEG studies targeting primary motor cortex link the N45 to the state of GABA_a_ receptor mediated inhibition and NMDA receptor mediated glutamatergic signalling [15, 16]. Extrapolating from these findings, our results indicate that early motor sequence learning is accompanied by a shift in E:I balance within SMA towards disinhibition.

A transient reduction in GABA inhibition supports learning by allowing an increase in NMDA receptor firing, and by extension, learning-related long-term potentiation [33]. To date, disinhibition associated with motor sequence learning has only been documented within M1 [3, 4, 10]. We extend these findings and show that disinhibition also occurs within higher-order motor cortex. We did not observe correlations between the N45 and learning and it might be the case that learning-induced disinhibition is important for aspects of learning we did not assess in the present study, such as consolidation over a longer timescale [26]. Although this possibility requires future investigation, our findings altogether imply that disinhibition is a neurophysiological mechanism within SMA that occurs during early learning of new motor sequences.

### The aperiodic exponent shifts across motor sequence learning and is associated with motor sequence improvements

The slope of 1/f exponential decrease in the EEG power spectral density is a recently established marker of E:I balance [17]. Contrary to expectations, and our N45 TEP results, we observed a gradual, non-significant increase in the aperiodic exponent (I > E) across learning, followed by a significant reduction post-learning (E > I). Notably, changes in exponents did not correlate with changes in TEPs, raising the possibility that these measures index different facets of neurotransmission. Pharmacological evidence suggests that the N45 and aperiodic exponent reflect both synaptic and extra-synaptic receptor signalling [15, 18, 34, 35], but the relative contribution of these pools of GABA to the N45 and aperiodic slope are currently unknown. Further research is thus required to examine the neurophysiological mechanism each marker reflects, and how each mechanism contributes to sequence learning. Nonetheless, our findings provide novel insights into the different E:I dynamics within SMA during sequence learning.

Interestingly, increases in the exponent across learning correlated with greater skill improvement. These correlations contrast with past findings of associations between lower cortical inhibition and better motor performance on different tasks [4, 12]. As the SVIPT is a complex motor sequence task [25], learning may have been bolstered by increased extra-synaptic inhibition which has been proposed to enhance the signal-to-noise ratio of information processing [36]. Altogether, these correlations suggest that links between the modulation of inhibition and motor learning are task and neural marker specific.

## Limitations

Most research into understanding the neurophysiological basis of TEP peaks comes from studies targeting M1. Current literature suggests the underlying neurotransmission of the N45 is site-specific [16, 37], and as such there is some uncertainty in interpreting the N45 evoked by SMA stimulation. Furthermore, our N100 findings should be interpreted with caution as control analyses revealed that the contribution of sensory vs. true effects of stimulation to this peak cannot be disentangled. Although the measures obtained in this study are time intensive, future research may include a control non-sequential motor condition (as in [4]) to examine the specificity of SMA disinhibition observed herein.

## Conclusion

Using TMS-EEG, we demonstrate disinhibition within SMA as participants learned novel motor sequences, and therefore provide new insight into the neurophysiological mechanisms of higher-order motor regions that accompany sequence learning. We also show that cortical inhibition indexed by the EEG aperiodic exponent is associated with learning outcomes. Future work is required to establish the synaptic or extra-synaptic nature of E:I dynamics that accompany motor sequence learning.

## Acknowledgements

We thank Jaeger Wongtrakun, Matthew Wiseman and Dr. Shou-Han Zhou for their assistance with data collection.

## References

1. Dayan, E. and L.G. Cohen, Neuroplasticity subserving motor skill learning. Neuron, 2011. 72(3): p. 443–54.

2. Ariani, G. and J. Diedrichsen, Sequence learning is driven by improvements in motor planning. J Neurophysiol, 2019. 121(6): p. 2088–2100.

3. Coxon, J.P., N.M. Peat, and W.D. Byblow, Primary motor cortex disinhibition during motor skill learning. J Neurophysiol, 2014. 112(1): p. 156–64.

4. Kolasinski, J., et al., The dynamics of cortical GABA in human motor learning. J Physiol, 2019. 597(1): p. 271–282.

5. Dupont-Hadwen, J., S. Bestmann, and C.J. Stagg, Motor training modulates intracortical inhibitory dynamics in motor cortex during movement preparation. Brain Stimul, 2019. 12(2): p. 300–308.

6. Nachev, P., C. Kennard, and M. Husain, Functional role of the supplementary and pre-supplementary motor areas. Nat Rev Neurosci, 2008. 9(11): p. 856–69.

7. Ashe, J., et al., Cortical control of motor sequences. Curr Opin Neurobiol, 2006. 16(2): p. 213–21.

8. Bachtiar, V. and C.J. Stagg, The role of inhibition in human motor cortical plasticity. Neuroscience, 2014. 278: p. 93–104.

9. Barron, H.C., Neural inhibition for continual learning and memory. Curr Opin Neurobiol, 2021. 67: p. 85–94.

10. Floyer-Lea, A., et al., Rapid modulation of GABA concentration in human sensorimotor cortex during motor learning. J Neurophysiol, 2006. 95(3): p. 1639–44.

11. King, B.R., et al., Baseline sensorimotor GABA levels shape neuroplastic processes induced by motor learning in older adults. Hum Brain Mapp, 2020. 41(13): p. 3680–3695.

12. Carlsen, A.N., J.S. Eagles, and C.D. MacKinnon, Transcranial direct current stimulation over the supplementary motor area modulates the preparatory activation level in the human motor system. Behav Brain Res, 2015. 279: p. 68–75.

13. Bortoletto, M., et al., The contribution of TMS-EEG coregistration in the exploration of the human cortical connectome. Neurosci Biobehav Rev, 2015. 49: p. 114–24.

14. Rogasch, N.C. and P.B. Fitzgerald, Assessing cortical network properties using TMS-EEG. Hum Brain Mapp, 2013. 34(7): p. 1652–69.

15. Premoli, I., et al., TMS-EEG signatures of GABAergic neurotransmission in the human cortex. J Neurosci, 2014. 34(16): p. 5603–12.

16. Belardinelli, P., et al., TMS-EEG signatures of glutamatergic neurotransmission in human cortex. Sci Rep, 2021. 11(1): p. 8159.

17. Donoghue, T., et al., Parameterizing neural power spectra into periodic and aperiodic components. Nat Neurosci, 2020. 23(12): p. 1655–1665.

18. Waschke, L., et al., Modality-specific tracking of attention and sensory statistics in the human electrophysiological spectral exponent. Elife, 2021. 10.

19. Gao, R., E.J. Peterson, and B. Voytek, Inferring synaptic excitation/inhibition balance from field potentials. Neuroimage, 2017. 158: p. 70–78.

20. Kaarre, O., et al., Association of the N100 TMS-evoked potential with attentional processes: A motor cortex TMS-EEG study. Brain Cogn, 2018. 122: p. 9–16.

21. Gordon, P.C., et al., Untangling TMS-EEG responses caused by TMS versus sensory input using optimized sham control and GABAergic challenge. J Physiol, 2023. 601(10): p. 1981–1998.

22. Biabani, M., et al., Characterizing and minimizing the contribution of sensory inputs to TMS-evoked potentials. Brain Stimul, 2019. 12(6): p. 1537–1552.

23. Oldfield, R.C., The assessment and analysis of handedness: the Edinburgh inventory. Neuropsychologia, 1971. 9(1): p. 97–113.

24. Cona, G., G. Marino, and C. Semenza, TMS of supplementary motor area (SMA) facilitates mental rotation performance: Evidence for sequence processing in SMA. Neuroimage, 2017. 146: p. 770–777.

25. Reis, J., et al., Noninvasive cortical stimulation enhances motor skill acquisition over multiple days through an effect on consolidation. Proc Natl Acad Sci U S A, 2009. 106(5): p. 1590–5.

26. Stavrinos, E.L. and J.P. Coxon, High-intensity Interval Exercise Promotes Motor Cortex Disinhibition and Early Motor Skill Consolidation. J Cogn Neurosci, 2017. 29(4): p. 593–604.

27. Rogasch, N.C., et al., Analysing concurrent transcranial magnetic stimulation and electroencephalographic data: A review and introduction to the open-source TESA software. Neuroimage, 2017. 147: p. 934–951.

28. Delorme, A. and S. Makeig, EEGLAB: an open source toolbox for analysis of single-trial EEG dynamics including independent component analysis. J Neurosci Methods, 2004. 134(1): p. 9–21.

29. Tadel, F., et al., Brainstorm: a user-friendly application for MEG/EEG analysis. Comput Intell Neurosci, 2011. 2011: p. 879716.

30. Team, J., JASP (Version 0.17.2.1) [Computer Software]. 2023.

31. Field, A., Miles, J., Field, Z., Discovering Statistics Using R. 2012: SAGE Publications Ltd.

32. Oostenveld, R., et al., FieldTrip: Open source software for advanced analysis of MEG, EEG, and invasive electrophysiological data. Comput Intell Neurosci, 2011. 2011: p. 156869.

33. Froemke, R.C., Plasticity of cortical excitatory-inhibitory balance. Annu Rev Neurosci, 2015. 38: p. 195–219.

34. Franks, N.P., General anaesthesia: from molecular targets to neuronal pathways of sleep and arousal. Nat Rev Neurosci, 2008. 9(5): p. 370–86.

35. Darmani, G., et al., Effects of the Selective alpha5-GABAAR Antagonist S44819 on Excitability in the Human Brain: A TMS-EMG and TMS-EEG Phase I Study. J Neurosci, 2016. 36(49): p. 12312–12320.

36. Koh, W., et al., GABA tone regulation and its cognitive functions in the brain. Nat Rev Neurosci, 2023. 24(9): p. 523–539.

37. Rogasch, N.C., et al., The effects of NMDA receptor blockade on TMS-evoked EEG potentials from prefrontal and parietal cortex. Sci Rep, 2020. 10(1): p. 3168.

