## Supplementary Data for "Supplementary motor area disinhibition during motor sequence learning: A TMS-EEG study"

### Supplementary Materials

### Supplementary Data 1

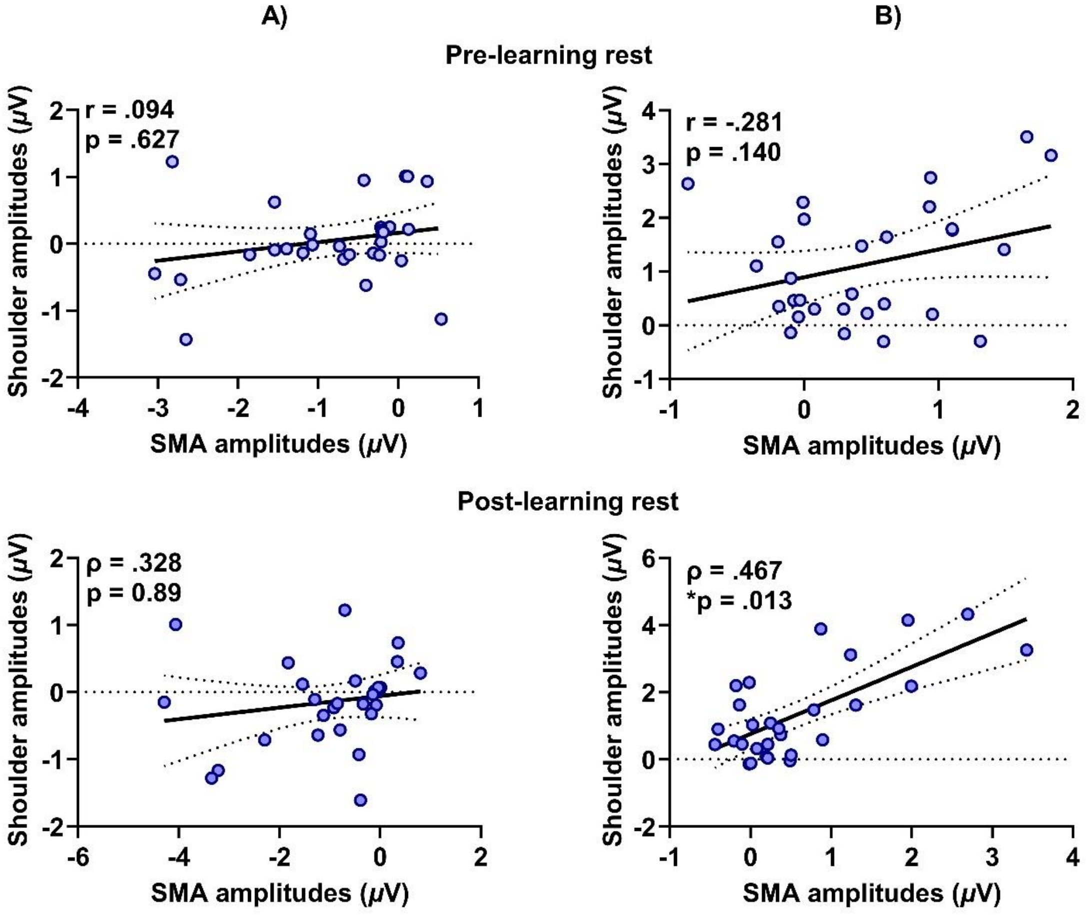


**Supplementary Figure 1.** Correlations between N45 **(A)** and N100 amplitudes **(B)** following SMA and shoulder stimulation at pre-learning rest (top panel) and post-learning rest (lower panel).

**Supplementary Data 2**

Greater (more negative) N100 amplitudes at baseline and early learning were associated with greater early and late skill (baseline: ρ = -.662, $p < .001$, early learning: ρ = -.462, $p = .012$) (baseline: ρ = -.520, $p = .004$, early learning: ρ = -.497, $p = .006$). No significant correlations were found between N100 amplitudes and delta skill (all *p*s > .7).

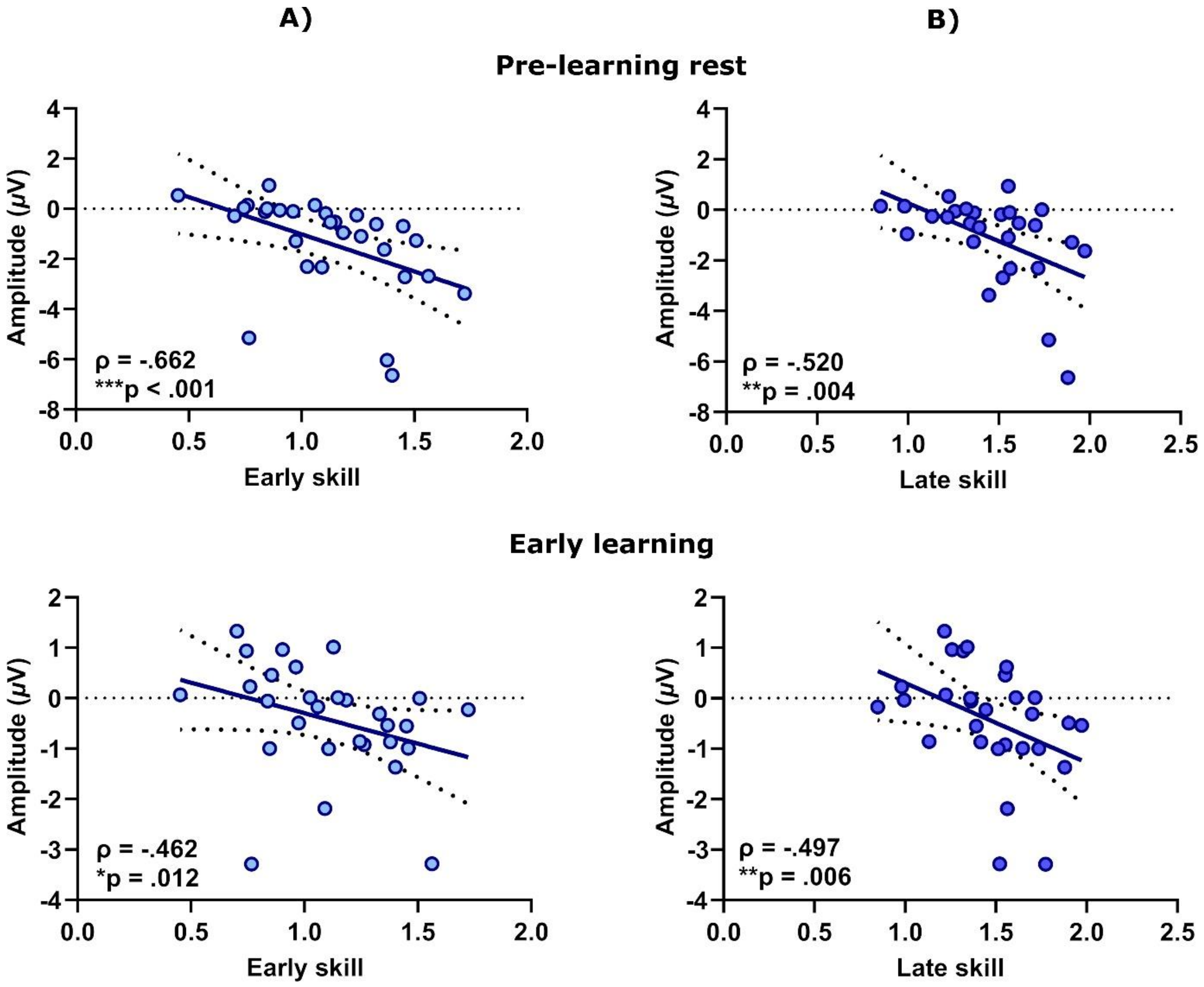


**Supplementary Figure 2.** Top panel: correlations between baseline N100 amplitudes and early skill **(A)** and late skill **(B).** Lower panel: correlations between early learning N100 amplitudes and early skill **(A)** and late skill **(B).**
